# TOWARDS CLIMATE-SMART REWILDING: AN INTEGRATED FRAMEWORK FOR BIODIVERSITY, CLIMATE CHANGE, AND SOCIETY

**DOI:** 10.1101/2025.03.21.644513

**Authors:** Gavin Stark, Magali Weissgerber, Néstor Fernández, Laura C. Quintero-Uribe, Marek Giergiczny, Nikolaj Rauff Poulsen, Nacho Villar, Bjorn Mols, Elisabeth S. Bakker, Angus Monro Smith, Georg Winkel, Diogo Alagador, José M. Rey-Benayas, Josep Maria Espelta, Miriam Selwyn, Lluís Brotons, Tatiana Kluvankova, Stanislava Brnkalakova, Judith Kloibhofer, Reinhard Prestele, Henrik G. Smith, Alba Lázaro-González, Robert Buitenwerf, Elena A. Pearce, Jens-Christian Svenning, Joana Santana, Pedro Beja, Francisco Moreira, Sven Wunder, Miroslav Svoboda, Vlado Vancura, Almut Arneth, Arndt Hampe, Henrique M. Pereira

**Affiliations:** German Centre for Integrative Biodiversity Research (iDiv) Halle-Jena-Leipzig, Leipzig, Germany; Institut für Biologie, Martin-Luther-University Halle-Wittenberg, Halle, Germany; 3Faculty of Economic Science, University of Warsaw, Warsaw, Poland; 4Department of Aquatic Ecology, Netherlands Institute of Ecology (NIOO-KNAW), Droevendaalsesteeg 10, 6708 PB Wageningen, Netherlands; Wildlife Ecology and Conservation Group, Wageningen University, P.O. Box 47 6700 AC Wageningen, The Netherlands; Forest and Nature Conservation Policy Group, Wageningen University, P.O. Box 47, 6700 AC Wageningen, the Netherlands; Biodiversity Chair, Mediterranean Institute for Agriculture, Environment and Development, MED/CHANGE, University of Évora, Évora, Portugal; University of Alcalá, Madrid, Spain; CREAF, Cerdanyola del Vallès 08193, Spain, Universitat Autònoma de Barcelona, E08193 Bellaterra (Cerdanyola del Vallès), Spain; CSIC, Cerdanyola del Vallès 08193, Spain; CTFC, Solsona 25280, Spain; SlovakGlobe, Forest Ecology Institute Slovak Academy of Sciences (SAS), Slovakia; Institute of Meteorology and Climate Research, Atmospheric Environmental Research (IMKIFU), Karlsruhe Institute of Technology (KIT), Garmisch-Partenkirchen, Germany; Department of Biology, Lund University, Sweden; INRAE, University of Bordeaux, BIOGECO, F-33610, Cestas, France; Center for Ecological Dynamics in a Novel Biosphere (ECONOVO), Department of Biology, Aarhus University, Ny Munkegade 114, DK-8000 Aarhus C, Denmark; BIOPOLIS Program in Genomics, Biodiversity and Land Planning, CIBIO, Campus de Vairão, 4485-661 Vairão, Portugal; CIBIO, Centro de Investigação em Biodiversidade e Recursos Genéticos, InBIO Laboratório Associado, Campus de Vairão, Universidade do Porto, 4485-661 Vairão, Portugal; CIBIO, Centro de Investigação em Biodiversidade e Recursos Genéticos, InBIO Laboratório Associado, Instituto Superior de Agronomia, Universidade de Lisboa, Tapada da Ajuda, Lisboa 1349-017, Portugal; European Forest Institute (EFI), St. Antoni M. Claret 167, 08025 Barcelona, Spain; Czech University of Life Sciences, Faculty of Forestry and Wood Sciences, Kamýcká 129, Praha 6 Suchdol, Czech Republic; Wilderness Director of European Wilderness Society and Wilderness Advocate, Austria

**Keywords:** climate change, ecosystem restoration, interdisciplinary, socio-economic, wildlife comeback

## Abstract

The UN Decade on Ecosystem Restoration and the Kunming-Montreal Global Biodiversity Framework aim to restore 30% of degraded ecosystems. Both the IPCC and IPBES highlight the crucial role of ecosystem restoration in addressing the interconnected crises of climate change and biodiversity loss. One key restoration strategy is rewilding, which enhances ecosystem complexity with minimal human intervention. While traditional rewilding strategies often focus on benefits for biodiversity, we propose a climate-smart rewilding framework as a new approach designed to deliver climate benefits alongside biodiversity restoration. This framework seeks to integrate biodiversity, climate change adaptation and mitigation, as well as socio-economic benefits and trade-offs. We illustrate how this framework can be utilized to identify areas across Europe where rewilding could increase carbon sequestration, enhance species’ abilities to adapt to the velocity of climate change, and maximize wildlife-watching benefits while minimizing the costs associated with livestock-wildlife conflict. Finally, we acknowledge some limitations of climate-smart rewilding, but we argue that its adaptability and low cost render it a promising solution to the challenges facing Europe and beyond.

## INTRODUCTION

We are currently amid the UN Decade on Ecosystem Restoration, during which the Kunming-Montreal Global Biodiversity Framework (CBD) aims to restore 30% of degraded ecosystems (CBD, 2022), a goal that is similarly echoed in other initiatives like the EU Biodiversity Strategy for 2030 (Hermoso et al., 2022) or the EU Restoration Regulation (Hering et al., 2023). While these legal frameworks prioritize nature conservation, there is a growing focus on climate change adaptation and mitigation, with policymakers increasingly seeking to utilize the carbon uptake and storage capacities of land ecosystems to help achieve climate change objectives (Bustamante et al., 2019; Cook-Patton et al., 2021; Griscom et al., 2017; von Holle et al., 2020). One effective approach to achieving these climate goals is through nature-based solutions (a concept coined by the World Bank; Sowińska-Świerkosz & García, 2022), which are seen as strategies to harnesses the power of nature to enhance carbon sequestration and bolster the resilience of ecosystems and local communities (Griscom et al., 2017; Marvin et al., 2023; Schmitz et al., 2023). Nature-based solutions have gained recognition in international discussions on climate change and biodiversity (Pörtner et al., 2023), with efforts focused on maintaining carbon sinks in vegetation and soil (Griscom et al., 2017). These initiatives include enhancing disaster risk reduction measures to mitigate the impacts of natural hazards, conserving native ecosystems, protecting forests, and restoring wetlands (Harenda et al., 2018; Lewis et al., 2019; McVittie et al., 2018; Nayak et al., 2022; Siikamäki et al., 2013).

In this context, rewilding emerges as a notable strategy for ecosystem restoration that is garnering increasing attention from both scientists and policymakers due to its potential benefits for biodiversity and society (Carver et al., 2021; Perino et al., 2019; Schmitz et al., 2023; Stark & Galetti, 2024; Stark & Schwarz, 2024; Svenning et al., 2016, 2024). Rewilding embraces a systems perspective that emphasizes the restoration of self-sustaining ecosystems (Perino et al., 2019). Rewilding has been proposed as a nature-based solution to address both biodiversity loss and climate change (Perino et al., 2019), although some core differences exist between the two approaches (see Waylen et al., 2024). By aiming to restore functional ecosystems, rewilding contributes to carbon storage and enhances ecosystems’ ability to withstand climate change for wildlife and society (Bakker & Svenning, 2018; Malhi et al., 2022; Pettorelli et al., 2019; Svenning, 2020; Villar & Medici, 2021). Furthermore, rewilding may offer many other benefits to people, including water purification and soil health (Cromsigt et al., 2018; Garrido et al., 2019; Harvey & Henshaw, 2023; Malhi et al., 2022; Sandom et al., 2020; zu Ermgassen et al., 2018).

Rewilding initiatives present notable opportunities to enhance socioeconomic outcomes (Faure et al., 2024). These initiatives have the potential to decrease management expenses and subsidies while fostering sustainable livelihoods in rural areas through ecotourism, the preservation of indigenous and local cultural heritage, and nature-based recreation (Schou et al., 2021; Williams et al., 2024). Additionally, they can benefit industries reliant on biodiversity, including agriculture (e.g., increasing pollinators, such as bees and butterflies), fisheries (e.g., restoring mangroves serves as nurseries for fish species), and forestry (increasing tree species diversity improves resilience against pests and diseases; Koninx, 2019; Massenberg et al., 2023; Pettorelli et al., 2019).

Climate change presents significant challenges to rewilding efforts by influencing the conditions under which they operate (Carroll & Noss, 2021; Malhi et al., 2016; Svenning et al., 2016). When undertaking rewilding initiatives, it is crucial to take into account a world characterized by species range shifts, community reshuffling, the disruption and formation of new biotic interactions, changing ecosystem dynamics, and an increasing frequency and intensity of stochastic disturbances (Bastazini et al., 2021; Boonman et al., 2022; Burak et al., 2024; Mokany et al., 2015). Such conditions may lead to alterations in species richness, community composition, and functional dynamics, as well as changes in biome distribution (Blois et al., 2013; Pecl et al., 2017). As species strive to maintain suitable environmental conditions and mitigate extinction risks, their ability to track climate shifts relies on the availability of natural corridors, underscoring the need for enhanced connectivity and secured dispersal routes (Sonntag & Fourcade, 2022). Climate change can also disrupt trophic processes, particularly when interacting species respond variably to changing conditions, especially in seasonal ecosystems with limited growth periods (Merz et al., 2023; Winder & Schindler, 2004). Additionally, climate change is expected to modify natural disturbance regimes, leading to intensified droughts, wildfires, floods, and other biotic disturbances (Jones et al., 2022; Seneviratne et al., 2021).

Currently, there is no formal framework that systematically integrates the influences of climate change on rewilding and the potential impacts of rewilding on climate-related consequences for ecosystems and livelihoods. To address this gap, we propose the concept of climate-smart rewilding, which enables a critical examination of the intricate connections among rewilding, biodiversity, climate change mitigation and adaptation, and socioeconomic factors that affect human well-being (Fig. 1). First, we present a comprehensive conceptual framework that outlines these interwoven relationships. Next, we explore practical examples based on this framework, demonstrating how various rewilding actions can effectively contribute to climate change mitigation, enhance the adaptive capacity of ecosystems, and deliver bundled biodiversity benefits for human well-being while minimizing associated social trade-offs, such as human-wildlife conflicts. Additionally, we illustrate how our framework can guide spatial prioritization of rewilding efforts to maximize socioecological benefits (for methodology, see supplementary materials). We conclude with a discussion of the climate-smart rewilding approach’s advantages and limitations.

**Figure 1.**
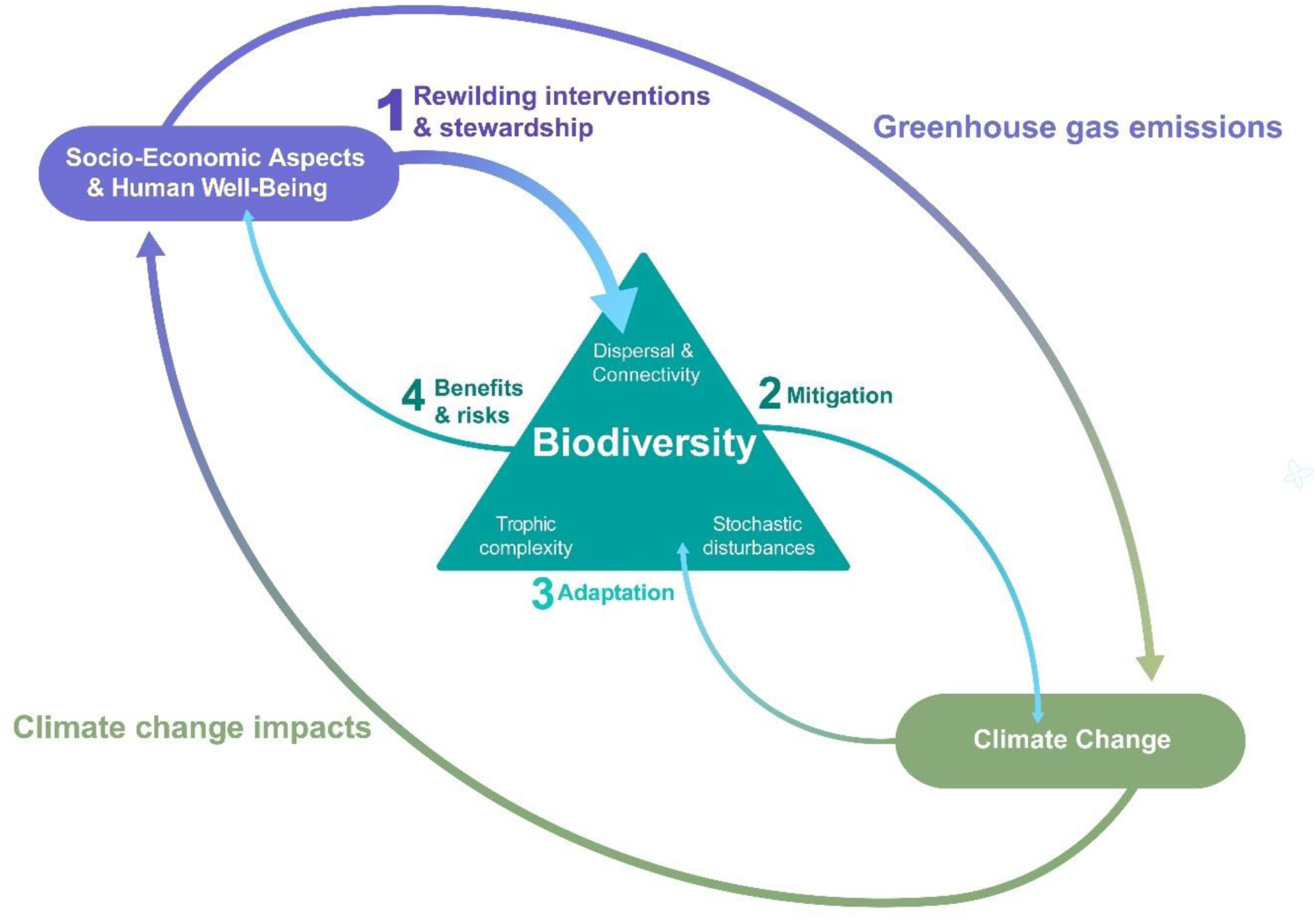
The new rewilding framework expands on the existing concept (Perino et al., 2019) by incorporating elements of climate change mitigation, adaptation, and rewilding impact on human society. It promotes climate-smart rewilding management actions (1) to restore ecological dynamics within rewilded ecosystems, encompassing targeted actions and unintended dynamics. This is achieved by enhancing the three components: dispersal and connectivity, trophic complexity, and stochastic disturbances. The result is an increase in biodiversity across ecosystems. The expansion of these rewilding components can lead to more dynamic and resilient ecosystems, contributing to climate change mitigation (2) and adaptation (3) efforts beneficial for both wildlife and society. The socio-economic dimension is also considered within the framework, aiming to cultivate sustainable strategies (by looking at the benefits and risks of rewilding) (4) that promote long-term stewardship of rewilded areas and encourage the development of more natural landscapes on a larger scale.

## CONCEPTUAL FRAMEWORK FOR REWILDING IN THE CONTEXT OF CLIMATE CHANGE

The climate-smart rewilding framework (Fig. 1) builds on and generalises the framework proposed by Perino et al. (2019), which highlights three key ecological processes as being the main target of rewilding: dispersal and connectivity, trophic complexity, and stochastic disturbances. The climate-smart rewilding framework places biodiversity at its core, implying that rewilding outcomes for climate change and human well-being will be mediated through its consequences for biodiversity. Compared to Perino et al. (2019), we here generalise the notion of biodiversity by adopting the three ecosystem level classes of Essential Biodiversity Variables (Fernández et al., 2020; Pereira et al., 2013), community composition, ecosystem structure, and ecosystem function, which have also been adopted by IPBES (IPBES, 2019). Within this generalised view, ecosystem structure may refer to landscape connectivity and how it mediates organisms’ dispersal and movement but also to structural vegetation complexity or microclimatic heterogeneity; community composition may refer to trophic interactions among biota (i.e., trophic complexity) but also to changes in the patterns of alpha and beta diversity; and ecosystem functioning may refer to stochastic disturbances but also to other functional aspects such as nutrient cycling. The generalised notion of biodiversity lends the Climate-Smart Rewilding framework a more explicit focus on ecosystem structure and functioning compared to Perino et al. (2019), and it should facilitate the establishment of functional links between rewilding outcomes and climate-change-driven dynamics of the abiotic and biotic environment.

The principal novelty of the climate-smart rewilding framework consists, however, in its systematic consideration of climate change mitigation and adaptation, both as drivers as well as outcomes of rewilding actions (see Fig. 1, processes 2 and 3). Mitigation and adaptation concern ecosystems not only directly, through their effects on biota and the abiotic environment, but also indirectly, through their implications for human societies and livelihoods. Therefore, socio-economic aspects and human well-being form a further integral part of the climate-smart rewilding framework. Socio-economic aspects can likewise be viewed both as drivers as well as outcomes of rewilding actions (Fig. 1, processes 1 and 4). Overall, we believe that an appropriate consideration and integration of all three framework components – biodiversity, climate change, and human well-being – will be crucial for successfully planning, conducting, and managing rewilding initiatives. The climate-smart rewilding framework thus advocates for rewilding strategies that maximize ecological, climate, and societal benefits while minimizing risks and trade-offs, thus promoting long-term sustainability, societal acceptance, and the resilience of natural landscapes (Table 1).

**Table 1.** Rewilding Intervention Management Decisions (a) and Their Consequences for Climate Change Adaptation and Mitigation (b), as well as the Benefits and Risks of Biodiversity to People (c). These are only examples. Several of these actions contribute to more than one ecosystem component, but we have allocated them to the main component they usually target.

| (a) | Trophic complexity (community composition and structure) | Connectivity (ecosystem spatial structure) | Stochastic disturbances (disruptions in ecosystem function) |
| --- | --- | --- | --- |
| **Rewilding actions** | • Creation of no hunting or fishing zones<sup>1</sup><br>• The reduction of wildlife-livestock conflicts through the adaptation of the husbandry system<sup>1</sup><br>• Facilitating the expansion of wildlife populations for instance by reducing domestic livestock grazing<sup>2</sup><br>• Reintroduction of native species with key functional roles<sup>3</sup> | • Removing dams, roads, fences, and any other man-made obstacles<sup>1</sup><br>• Manage human access to recorded and predicted migration routes for plants and animals, including birds and insects, to create landscapes that are more permeable to movement<sup>1,4</sup><br>• Repurpose abandoned landscapes for habitat connectivity<sup>4</sup> | • Reduce human activities that appropriate net primary productivity, for instance by reducing forest harvest, livestock grazing, or agricultural interventions<sup>1</sup><br>• Occasionally reduce the suppression of wildfires ignited within rewilded areas<sup>1</sup><br>• Restore natural hydrological regimes and land-water interactions<sup>5</sup><br>• Reduce carrion and deadwood removal and supplemental feeding activities<sup>1</sup> |
| (b) |  |  |  |
| **Climate Change mitigation** | • Replacing domestic ruminants with large wild non-ruminant herbivorous species can decrease methane (CH<sub>4</sub>) emissions<sup>6</sup><br>• More natural grazing by wildlife reduces fire frequency and overall carbon emissions<sup>7</sup><br>• Trampling by grazers/browsers can increase soil carbon storage/reduce C soil emissions<sup>8</sup><br>• Seed dispersal of large-seeded trees by large birds and scatter hoarders can enhance C sequestration per unit tree<sup>9</sup> | • Increasing long-distance seed dispersal allows the expansion of forests, grasslands, and other vegetation forms, enhancing the potential for carbon uptake and storage in these ecosystems i.e., via allowing plant species to track climate change<sup>9</sup><br>• More connected diversity of habitats increases vegetation complexity and carbon stocks<sup>10</sup><br>• Increasing net primary productivity in connected abandoned landscapes<sup>11</sup> | • A more natural rate of stochastic disturbances (and increasing active management to avoid fire propagation from outside the rewilded area) can decrease the amount of carbon and other GHG that are released into the atmosphere<sup>12</sup><br>• Restoring flooding regimes can shift the role of riparian zones from a carbon source to a carbon sink<sup>13</sup><br>• The expansion of wetlands can significantly enhance long-term carbon sequestration, capturing and storing substantial amounts of carbon from the atmosphere<sup>14</sup> |
| **Climate Change Adaptation** | • The presence of key ecosystem engineers (e.g., beavers) increases heterogeneity and complexity inside an ecosystem, increasing its resilience to climate change<sup>15</sup> | • Connecting different ecosystems can increase access to potentially more areas with thermal refugia, vegetation cover, and access to water resources in different | • Permitting the occurrence of disturbances, such as natural wildfires in rewilded areas, can foster a mosaic of vegetative stages that enhance biodiversity and subsequently |
|  | Increasing vegetative habitat diversity due to natural grazing can increase potential refuges for different species under increasing temperatures<sup>16</sup><br>Reintroducing a combination of grazers and browsers can control woods expansion, generating more complex mosaics (e.g., grass and shrublands) of habitats that are resistant to extreme climatic events (e.g., limiting fire spread)<sup>17</sup> | seasons, reducing the impact of extreme climate events (e.g., heat waves)<sup>18</sup><br>Increasing intact migration routes and stepping stones can allow animals to reduce their exposure to climate fluctuations and rapid phenological timing changes while also being important for plant dispersal (and hence vegetation) mediated via the large herbivores<sup>19</sup><br>Increasing dispersal can improve gene flow from one habitat patch to another, improving the functional adaptability of populations to rapid climatic changes<sup>20</sup> | improve the resilience of ecosystems facing temperature fluctuations associated with climate change<sup>21</sup><br>Restoring natural river flow can help migrating aquatic species to reach areas where climate fluctuations are more stable<sup>22</sup><br>Increasing adaptability and resilience through more diversity and heterogeneity by reducing management in forests and other vegetation<sup>23</sup> |
| (c) |  |  |  |
| **Socio-economic Benefits** | Increasing opportunities for enjoyment of biodiversity and wildness contributing to psychological and physical health<sup>24</sup><br>Fostering social cohesion and enhancing a sense of identity and pride related to wildlife among local communities that preserve cultural heritage<sup>25</sup><br>Enabling sustainable ecotourism that benefits both local inhabitants and visitors<sup>26</sup> | More connectivity between urban and natural areas, can increase access to wilderness and provide myriad advantages to people (e.g., recreation, bike and walking routes with natural surroundings)<sup>27</sup><br>Wider access to water sources among different communities along the restored rivers<sup>28</sup><br>Increased connectivity among natural habitats can enhance ecosystem services such as pollination and pest control, which are vital for agricultural productivity. This can lead to more stable yields and reduced reliance on chemical inputs<sup>29</sup> | Increasing regulation of water quality & quantity for urban or rural areas<sup>29</sup><br>Disaster risk reduction of floods<sup>30</sup><br>Restoring ecosystems helps to provide quality environments and puts urban people in closer contact with nature, while in rural areas it may also provide sustenance<sup>31</sup><br>Deadwood can increase biodiversity and recreational choices for forest visitors<sup>32</sup> |
| **Socio-economic risks** | Damages to crops and livestock<sup>33</sup><br>Disease spreading from interacting with wild animals<sup>33</sup><br>Economic burden of wildlife comeback (e.g., reintroduction program expenses, increasing food prices, car crashes with wildlife)<sup>33</sup> | Increased risks of interactions between people and animals (human-wildlife conflicts) due to more natural areas in proximity to urban regions<sup>33</sup><br>Reduction in land available for agriculture, forestry, or urban development<sup>34</sup><br>Increased risk of frequent (due to climate change) extreme | Increased exposure to natural wildfires (from rewilded areas)<sup>36</sup><br>Increased flooding (due to decreasing flood management) in adjacent to rural or urban areas<sup>37</sup><br>Increased risk of forest fires and insect infestation to decreased recreational |
|  |  | events (mega-fires, floods,<br>etc.) reaching urban areas<sup>35</sup> | opportunities and site<br>attractiveness<sup>38</sup> |

The full reference list follows the numbers for each rewilding action and is presented following the general list in the
references section.

Given climate change’s potential to disrupt ecosystems (Carroll & Noss, 2021; Svenning et al., 2024), carefully selected implementation actions are crucial (Bergin et al., 2024). We reviewed the literature to identify interventions targeting the three key ecosystem processes addressed by rewilding (see Table 1). Additionally, we examined how rewilding interventions focusing on these dimensions of biodiversity and their synergistic effects can contribute to climate mitigation, climate adaptation, and socio-economic benefits and risks. For example, restoring ecological corridors on abandoned farmland—such as allowing natural vegetation to regenerate along former field margins—facilitates the movement and range expansion of species, enabling them to track climate change (Pereira & Navarro, 2015; Table 1a). In the face of rapid climate change, landscapes characterized by complex and well-connected ecosystems provide improved microclimatic buffering and access to climate refugia (Table 1b; Morelli et al., 2016; Stark et al., 2023). Another key intervention is the reintroduction of native grazers and browsers, which play critical ecological roles in enhancing biodiversity while also contributing to the reduction of methane emissions from livestock and promoting carbon sequestration through more varied grazing patterns (Holdo et al., 2009; Wilson & Edwards, 2008; Table 1b). Moreover, facilitating the expansion of wildlife populations can support sustainable ecotourism, benefiting both local communities and visitors. However, it is important to consider the potential for increased human-wildlife conflicts (Faure et al., 2024; Table 1c).

Selecting the best rewilding strategy involves balancing socio-economic trade-offs while focusing on approaches that increase resilience to climate change and benefit biodiversity. For the climate-smart Rewilding framework to gain practical implementation and raise support from policymakers, its fundamental elements must encompass the socio-economic impact of rewilding actions on society and human well-being. The impact of rewilding management on society can be evaluated by assessing the benefits and costs associated with restoring biodiversity and increased interactions between people and natural areas (Fig. 1 and Table 1). The holistic approach to rewilding, which promotes ecological integrity (Torres et al., 2018) alongside reducing the impacts of climate change on nature and people, would be highly recommended (Selwyn et al., 2025).

## APPLYING THE FRAMEWORK: THREE EXAMPLES

The following examples illustrate the application of the climate-smart rewilding framework to identify areas in Europe where rewilding actions can generate socio-economic and climate-related benefits. It is important to note that the spatial analyses presented here are illustrative, demonstrating the framework’s utility rather than providing definitive, Europe-wide prioritization maps. Instead, these examples highlight the diverse benefits and trade-offs associated with rewilding, emphasizing the need for context-specific assessments and the integration of social and economic considerations into rewilding strategies. They are intended to stimulate discussion around the potential of climate-smart rewilding in various European settings and encourage more nuanced, localized analyses.

## EXAMPLE 1: CLIMATE CHANGE MITIGATION ON ABANDONED FARMLANDS

The European Union has set up ambitious goals for restoring degraded ecosystems and enhancing their capacity to take up and store carbon dioxide, yet it remains an object of debate how these goals can be realistically achieved. Rewilding abandoned farmlands could be one highly feasible and cost-effective approach (Pereira & Navarro, 2015). Farmland abandonment is a widespread phenomenon in many European regions (Keenleyside & Tucker, 2010), as a result of the migration of rural residents to urban areas in search of better opportunities, inadequate infrastructure, the remoteness of regional centers, difficulties in land management, low soil fertility combined with insufficient funds for improvement, and the reduced labor demands resulting from advancements in agricultural technologies (Levers et al., 2018; Suziedelyte Visockiene et al., 2019). Projections estimate that around 200,000 km^2^ of EU farmlands are at high risk of abandonment over the coming years (Castillo et al., 2018). Where no competing land uses exist, rewilding can be readily implemented as a low-impact management strategy that recreates self-sustaining ecosystems while efficiently increasing carbon stocks.

Applying our climate-smart Rewilding framework (Fig. 1), we identified regions in Europe that are particularly suited for carbon storage through passive rewilding (a non-interventionist approach), taking into account societal opportunity costs, such as farmland use. To evaluate the first component, we mapped potential net primary productivity (NPP; Plutzar et al., 2016). For the second component, we assessed the estimated probability of farmland abandonment (detailed in Supplementary Materials A regarding our spatial analysis and data collection). By integrating these two types of spatial data, we can identify areas where implementing passive rewilding as a management strategy could most effectively contribute to climate change mitigation while also considering competing land uses. Higher NPP, assuming all other factors are equal, indicates greater potential for increased standing biomass and aboveground carbon sequestration. Our spatial analysis reveals that abandoned agricultural landscapes are concentrated in southern and eastern Europe (Fig. 2A, Palmero-Iniesta et al., 2021). We also identified potential restoration areas with high net primary productivity (NPP) in the central-eastern regions (Fig. 2A). Overlaying the maps of predicted agricultural land abandonment and potential NPP highlights key areas for rewilding that could significantly enhance carbon sequestration, including the south of France, northern Spain, Portugal, and various parts of Eastern Europe (Fig. 2A).

**Figure 2.**
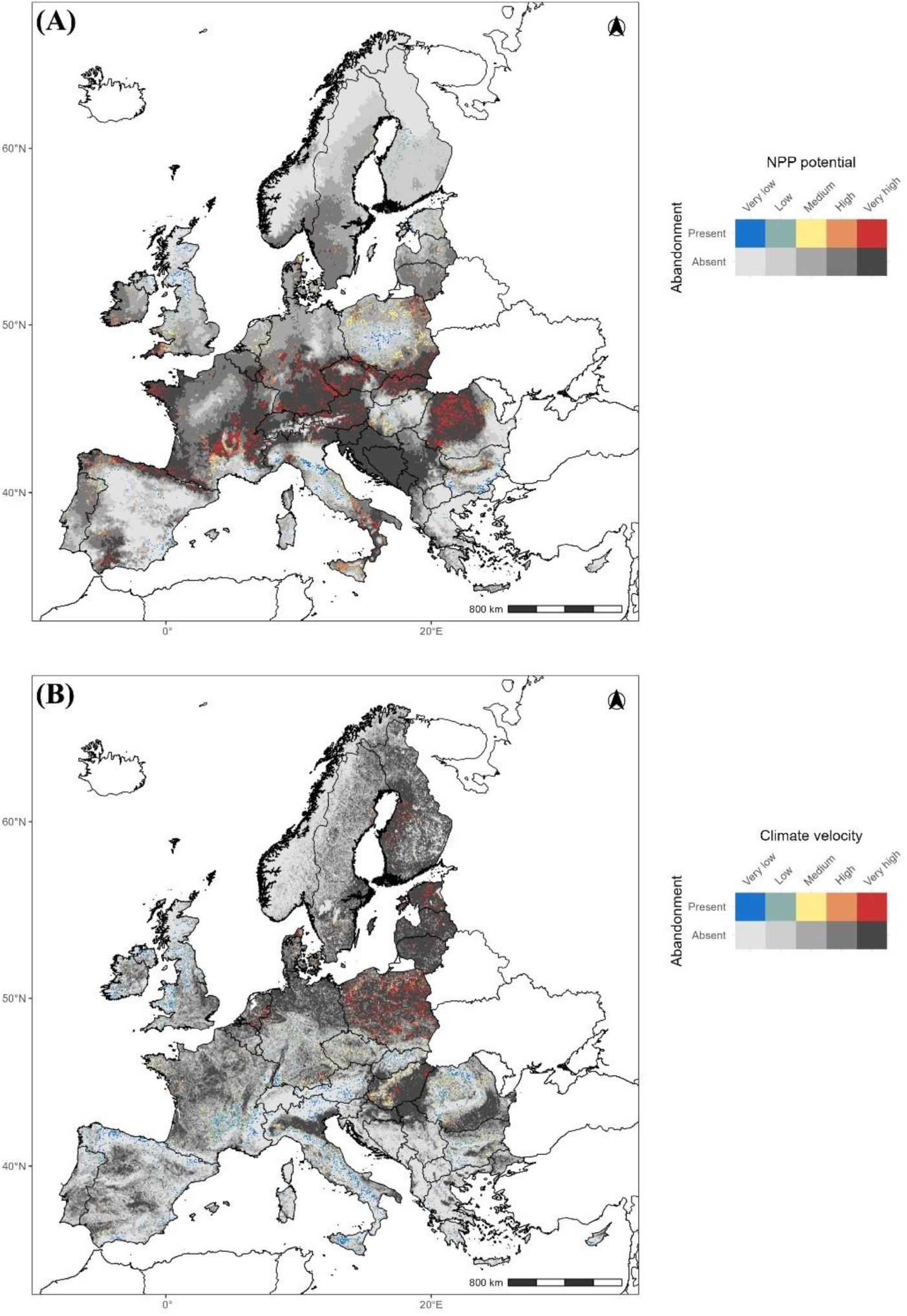
Spatial analysis of potential European rewilding areas for climate change mitigation (A) and adaptation (B) using a climate-smart framework based on abandoned landscapes (Ceaușu et al., 2015) and current land use (for more details refer to supplementary materials). (A) shows areas suitable for enhanced vegetative biomass (net primary productivity, NPP; blue to red: low abandonment probability/NPP to high abandonment probability/NPP) and carbon sequestration. (B) identifies potential ecological corridors facilitating species’ climate change tracking by mapping climate velocity over the next century.

Previous studies have indicated that the expansion of vegetation is often accompanied by an increased risk of large wildfires, particularly within Mediterranean regions (García-Ruiz et al., 2020; Vieira et al., 2023). To solve this, managing fires to align with natural fire regimes can reduce their frequency compared to current conditions, thereby lowering CO2 emissions (Jones et al., 2022; Rabin et al., 2022). Moreover, in disturbed environments, this is often followed by the expansion of invasive plant species that become dominant, resulting in simplified and species-poor landscapes with an increased risk of wildfires (Suárez-Ronay et al., 2024). This phenomenon poses a potential threat to the carbon sequestration benefits that are typically associated with rewilding initiatives aimed at climate change mitigation. Therefore, it is imperative to incorporate a variety of rewilding management strategies (or other land use management strategies that increase carbon intake, such as active reforestation; Di Sacco et al., 2021) that can generate synergistic outcomes. This can involve the introduction of low-intensity fires, reintroducing native wild grazers and browsers, protecting natural migration routes for wildlife, and allowing limited and managed grazing by domestic animals (Table 1), all aimed at reducing fuel loads and hence reducing large wildfire risk (Cromsigt et al., 2018). Identifying areas suitable for strategic restoration, including spontaneous tree regeneration alongside reintroducing native herbivores, is essential (Fuhlendorf et al., 2009; Malhi et al., 2022; Pearce et al., 2025). This assessment adopts a “supply-side” perspective, emphasizing the availability of land with lower barriers towards rewilding and the capacity for effective carbon sequestration.

## EXAMPLE 2: CLIMATE CHANGE ADAPTATION AND HAZARD PREVENTION

Rewilding strengthens the natural capacity of ecosystems to adapt to climate change (Svenning et al., 2024) by enhancing connectivity and fostering the creation of ecological corridors (Perino et al., 2019). These corridors are essential for linking fragmented habitats, allowing for greater species movement and genetic exchange (Carver et al., 2021). This connectivity is essential for maintaining stable populations and ensuring that ecosystems can adapt to shifting environmental conditions (Marjakangas et al., 2018; Mittelman et al., 2022). Rising temperatures disrupt ecosystems and drive species to adjust behaviors, physiology, migration patterns, and the timing of life cycle events (Charmantier et al., 2008; Thakur et al., 2020). While some species relocate to cooler or more favorable habitats, others modify their breeding, phenology, or hibernation schedules to align with changing climate patterns (Hoffmann & Sgrò, 2011). Those species unable to adapt face heightened extinction risks, potentially leading to biodiversity loss (Radchuk et al., 2019).

The restoration of ecosystems also plays a critical role in enhancing society’s resilience to climate impacts, which can mitigate the adverse impacts of extreme events (Munang et al., 2013; Scarano, 2017). Restored ecosystems can buffer against climate-induced hazards such as mega-fires, floods, droughts, and extreme weather events (Munang et al., 2013; Paudel et al., 2024; Scarano, 2017). For example, rewilding wetlands can potentially absorb excess water during storms and prevent flooding while maintaining water availability during droughts (Harvey et al., 2024). Moreover, restoring forests and grasslands enhances soil quality, preventing erosion and maintaining water retention, which can improve agricultural productivity and food security during droughts (Lal, 2015). Thus, diverse ecosystems are better able to adapt to changing conditions, ensuring the continued provision of essential services such as clean water, food, and shelter for both humans and wildlife (Munang et al., 2013).

Applying our climate-smart rewilding framework, we performed a spatial exploration of areas that could serve as migration routes or potential habitats for animal species to reoccupy (which could indirectly help in plant species migration via seed dispersal), thereby helping to mitigate the impact of unstable climates in the future. We begin by pinpointing regions that facilitate wildlife connectivity, characterized by minimal competing land uses and subsequent ecosystem changes, particularly concentrating on previously evaluated abandoned areas intended for climate change mitigation. We then analyze the climate stability of these regions by assessing the predicted rate of temperature change, known as climate velocity, to determine their potential as rewilding areas (for more details, see Supplementary Materials A). In general, species inhabiting high-velocity areas will need to move more swiftly to track their climatic niche than in low-velocity areas (Burrows et al., 2014). Therefore, the restoration of large-scale corridors is most essential in high-climate-velocity areas. This European-scale analysis aimed to assess whether these areas could serve as pathways for species migrating from areas experiencing rapid climate change to more stable ones. According to our spatial analysis, this phenomenon is especially noticeable in the eastern parts of the continent, notably in countries such as Poland, the Baltic States, and Slovakia (Fig. 2B). In contrast, several regions in southern Europe—such as northern Portugal, Spain, and the Italian peninsula—exhibit low climate velocity and a significant amount of abandoned land. This emphasizes the importance of prioritizing these areas for protection, as the need for large corridors to facilitate species range shifts is less urgent here than in other regions (Fig. 2B). Transforming abandoned lands into dynamic mosaic landscapes with varied vegetation stages, including forests, grasslands, and shrublands, may help link critical habitats across Europe and encourage animal movement towards more stable climates.

## EXAMPLE 3: SOCIAL BENEFITS AND TRADE-OFFS RELATED WITH WILDLIFE COMEBACK

Rewilding provides both opportunities and challenges for society. On one hand, the potential societal benefits of climate-smart rewilding are extensive (Table 1c). Rewilding degraded landscapes into healthy ecosystems supports human well-being by providing clean air, water filtration, food security, disease regulation, and resilience to natural hazards, such as flood or natural wildfire regulation (Bennett et al., 2015; Summers et al., 2012). Rewilding can also promote ecotourism, sustainable land use, and new economic opportunities for local communities while reducing subsidies for ecologically damaging activities and providing alternatives to traditional or intensive land-use practices (Hall, 2019; Pellis, 2019; Thulin & Röcklinsberg, 2020). In rural areas affected by environmental degradation, rewilding can revive economic prospects and enhance social resilience (Burnet et al., 2021; Schou et al., 2021). In certain circumstances, restored natural environments may offer important psychological and health benefits, enhancing mental well-being and reducing stress, especially in urban areas (Bratman et al., 2019). Finally, studies in different countries point to the preference of at least substantial parts of the European population for elements corresponding with “wilder” landscapes, such as rich-structured forest landscapes (Dunn-Capper et al., 2024; Giergiczny et al., 2015).

We examine regions where natural expansions increase trophic complexity to reveal important ‘demand-side’ impacts on human society. We illustrate this with two cases: The first showcases the positive effects of wildlife observation (Browning et al., 2024), while the second addresses a specific instance of human-wildlife conflict (Pimenta et al., 2017, 2018). In this context, the social benefits of rewilding are estimated using the concept of willingness to travel (WTT), as described by Giergiczny et al. (2022). WTT reflects the distance individuals are willing to travel to observe specific animal species. We integrated this data with human population density and overlaid the results with potential wildlife presence (see Supplementary Materials B for more details). In addressing the risks of rewilding to society, particularly regarding the potential for increased human-wildlife conflicts associated with increasing trophic complexity, we focused on identifying regions where wildlife may pose a threat to livestock, primarily through predation. Our objective was to highlight areas where the potential expansion of large predators in Europe coincides with varying densities of cattle, sheep, and goats (Gilbert et al., 2018).

Our study has pinpointed specific regions where the return of certain species (specifically, eight iconic species of large herbivores and carnivores; for further details, refer to the supplementary materials B) is anticipated. Geographical regions that are most likely to see a resurgence in wildlife, which may encourage people to undertake longer journeys, are primarily located in many mountainous areas across Europe. This includes the northern part of the Iberian Peninsula (particularly around the Pyrenees), the Alps, the Balkans, and various areas in Eastern Europe (Fig. 3A). This trend aligns with the potential economic benefits that ecotourism could bring to these areas, although it is important to note that ecotourism also depends on the aesthetic value of the landscapes. In addressing the risks of rewilding to society, regions most susceptible to the risks associated with rewilding and heightened human-wildlife conflicts include the Balkans, Western Carpathians, the Apennines in Italy, and parts of northern Portugal and Spain, particularly around Galicia and Asturias (Fig. 3B).

**Figure 3.**
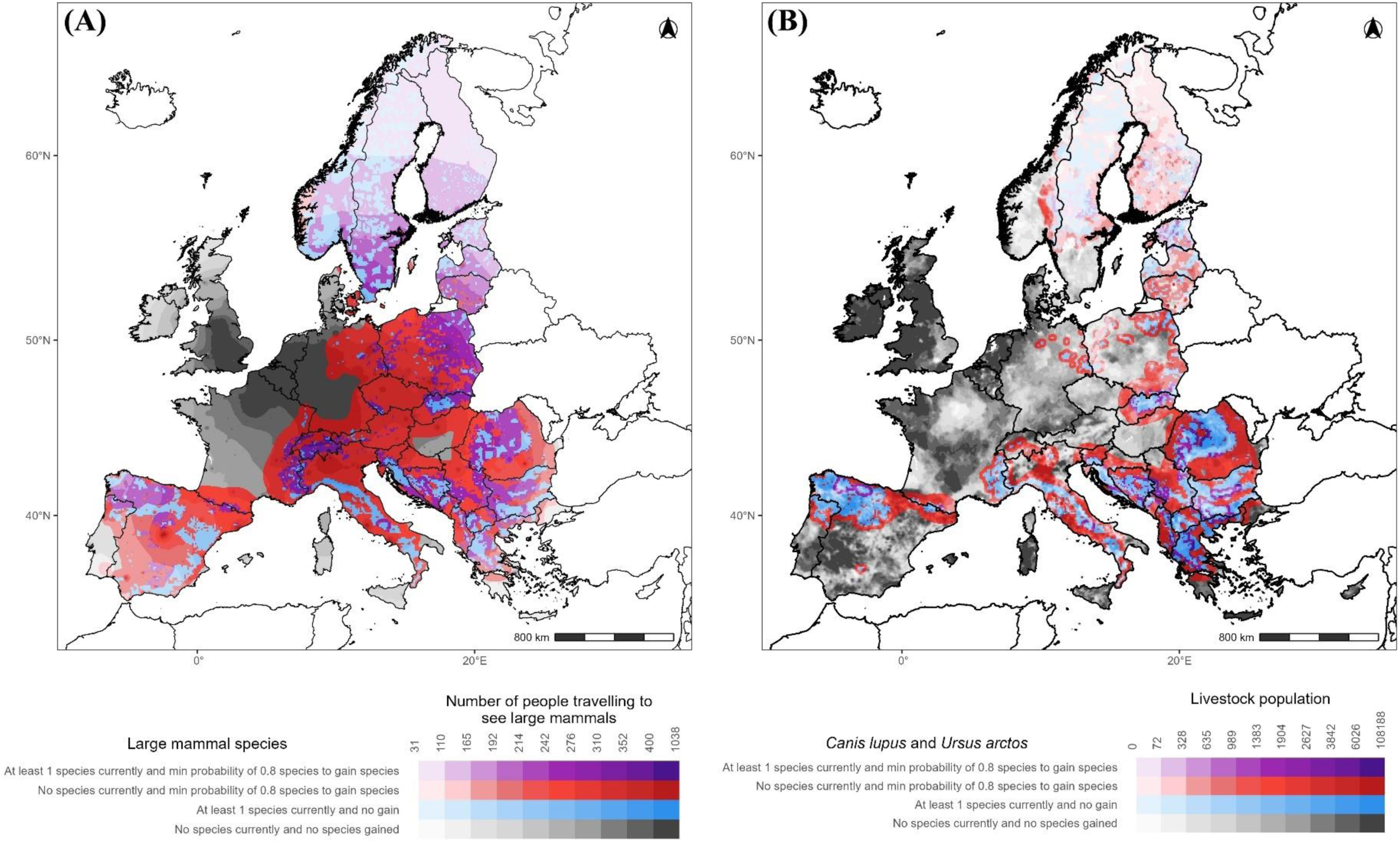
Spatial distribution of potential social benefits (A) and risks (B) associated with rewilding. (A) displays an index that combines the likelihood of colonization/expansion of eight large mammal species with visitor numbers (willingness to travel). (B) presents an index that factors in the likelihood of colonization by *Canis lupus* and *Ursus arctos* alongside livestock density. The analysis focuses only on regions expected to gain new species, excluding areas with pre-existing species. Color scheme: Purple – at least one species present and gained; Red – no species present and at least one species gained; Blue – species present with no gain; Black – no species present or gained.

Our spatially explicit maps highlight potential trade-offs between recreational benefits and local livelihoods in the Northern Iberian Peninsula, various parts of the Balkans, and regions of Eastern Europe, including Romania, Hungary, and Southern Poland (Fig. 3). In places like the Iberian Peninsula, some measures are already being implemented to mitigate these conflicts, such as modifying husbandry practices by fencing cattle at night during winter. This approach can substantially reduce predation, even in areas with significant livestock-predator overlap; however, it may not be universally effective (Álvares et al., 2014).

## CONCLUSIONS AND OUTLOOK

This study introduces a novel framework aimed at guiding rewilding initiatives that not only enhance biodiversity and societal outcomes but also effectively address the challenges posed by global climate change. This climate-smart rewilding framework builds upon earlier methodologies by integrating aspects of climate change mitigation and adaptation alongside socio-economic considerations into the planning and execution of rewilding efforts. By acknowledging the critical biodiversity-climate nexus, our approach results in a more comprehensive and policy-relevant strategy. The main objective of this framework is to prioritize species adaptation and highlight the ecosystem’s role in carbon sequestration, ultimately promoting self-sustaining ecosystems as integral components of rewilding design. Furthermore, embedding rewilding initiatives within climate change mitigation efforts and thoroughly evaluating their socio-economic impacts will be vital for policymakers and conservationists as they seek effective strategies to combat biodiversity loss and address climate challenges (Root-Bernstein et al., 2018).

Climate-smart rewilding marks a transformative shift in our approach to the interlinked crises of climate change and biodiversity loss. By harnessing nature’s potential for both mitigation and adaptation, it offers pathways for ecosystem restoration and societal benefits. Effective implementation, however, demands meticulous planning, community involvement, and adaptive management to address potential risks and challenges. It is important to note that while climate change often shifts focus toward immediate mitigation and disaster response (Pettorelli et al., 2019), there are also unique opportunities arising from it. For example, as the productivity of agricultural lands declines, previously cultivated areas present valuable prospects for conservation and ecosystem restoration (Bell et al., 2020; Pereira & Navarro, 2015; Rey Benayas et al., 2007).

While the framework presents numerous promising avenues, it is essential to acknowledge its limitations, particularly concerning spatial analysis, which may oversimplify the intricate interactions between biodiversity, climate change, and socio-economic factors. This simplification can lead to varied assessments of economic benefits and risks, including potential conflicts between human populations and wildlife. Additionally, the effectiveness of the framework can vary with context, necessitating modifications to suit different scales and ecological conditions. Therefore, establishing clear success criteria and measurable indicators for climate-smart rewilding is crucial for its effective application. This framework and its spatial analysis also shine a light on potential trade-offs within three key ecological components of rewilding: trophic complexity, connectivity, and stochastic disturbances, particularly in the context of climate change mitigation and adaptation. For example, enhancing habitat connectivity through large-scale restoration can support species movement but may elevate wildfire risks, potentially compromising carbon sequestration efforts. Likewise, re-establishing natural disturbance regimes might improve ecosystem integrity while posing challenges to local livelihoods. Such scenarios underscore the necessity for tailored strategies that thoughtfully assess these trade-offs. A standardized, one-size-fits-all model is insufficient; optimal intervention strategies will vary significantly based on localized ecological conditions, climate variability, socio-economic contexts, and specific rewilding goals—whether they involve carbon sequestration, biodiversity recovery, or resilience enhancement.

Looking ahead, future research should aim to explore a diverse range of intervention strategies beyond those presented in Table 1, refining local selection criteria to optimize rewilding outcomes (Torres et al., 2018). This exploration could include methods to boost trophic complexity, detailed analyses of connectivity to evaluate corridor designs, and modeling of disturbance regimes to balance ecosystem resilience with socio-economic aims. To ensure the broader applicability of these strategies, adapting indicators to regional contexts, integrating indigenous knowledge systems, and tackling diverse governance challenges will be vital (Lam et al., 2020). Comprehensive case studies across varied biomes will further validate the framework’s effectiveness (Convery et al., 2025). In conclusion, successfully implementing climate-smart rewilding strategies requires the incorporation of adaptive management frameworks, robust monitoring and evaluation methods, and investments in capacity-building and knowledge-sharing initiatives. This framework provides valuable guidance for policy and restoration efforts, aligning with the Convention on Biological Diversity (CBD) and EU biodiversity restoration targets (Friedman et al., 2022; Hering et al., 2023). By contributing to these important goals, our approach can mobilize additional resources and drive meaningful progress in addressing the intertwined challenges of climate change and biodiversity loss.

## Acknowledgments

We are grateful to Jeremy Dertien, Marta Cimatti, Luise Quoss, Matthias Baumann, and Tobias Kuemmerle for insightful discussions and data sharing during this study’s design.

## References for Table 1

1) Torres, A., Fernández, N., Zu Ermgassen, S., Helmer, W., Revilla, E., Saavedra, D., … & Pereira, H. M. (2018). Measuring rewilding progress. Philosophical Transactions of the Royal Society B: Biological Sciences, 373(1761), 20170433.

2) Perino, A., Pereira, H. M., Navarro, L. M., Fernández, N., Bullock, J. M., Ceaușu, S., … & Wheeler, H. C. (2019). Rewilding complex ecosystems. Science, 364(6438), eaav5570.

3) Svenning, J. C., Pedersen, P. B., Donlan, C. J., Ejrnæs, R., Faurby, S., Galetti, M., … & Vera, F. W. (2016). Science for a wilder Anthropocene: Synthesis and future directions for trophic rewilding research. Proceedings of the National Academy of Sciences, 113(4), 898-906.

4) Navarro, L. M., & Pereira, H. M. (2015). Rewilding abandoned landscapes in Europe. In Rewilding European landscapes (pp. 3-23). Cham: Springer International Publishing.

5) Harvey, G. L., & Henshaw, A. J. (2023). Rewilding and the water cycle. Wiley Interdisciplinary Reviews: Water, 10(6), e1686.

6) Wilson, G. R., & Edwards, M. J. (2008). Native wildlife on rangelands to minimize methane and produce lower-emission meat: kangaroos versus livestock. Conservation Letters, 1(3), 119-128.

7) Malhi, Y., Lander, T., le Roux, E., Stevens, N., Macias-Fauria, M., Wedding, L., … & Canney, S. (2022). The role of large wild animals in climate change mitigation and adaptation. Current Biology, 32(4), R181-R196.

8) Kaštovská, E., Mastný, J., & Konvička, M. (2024). Rewilding by large ungulates contributes to organic carbon storage in soils. Journal of Environmental Management, 355, 120430.

9) Bello, C., Crowther, T. W., Ramos, D. L., Morán-López, T., Pizo, M. A., & Dent, D. H. (2024). Frugivores enhance potential carbon recovery in fragmented landscapes. Nature Climate Change, 1-8.

10) Lundberg, J., & Moberg, F. (2003). Mobile link organisms and ecosystem functioning: implications for ecosystem resilience and management. Ecosystems, 6, 0087-0098.

11) Gvein, M. H., Hu, X., Næss, J. S., Watanabe, M. D., Cavalett, O., Malbranque, M., … & Cherubini, F. (2023). Potential of land-based climate change mitigation strategies on abandoned cropland. Communications Earth & Environment, 4(1), 39.

12) Linley, G. D., Jolly, C. J., Doherty, T. S., Geary, W. L., Armenteras, D., Belcher, C. M., … & Nimmo, D. G. (2022). What do you mean,‘megafire’?. Global Ecology and Biogeography, 31(10), 1906-1922.

13) Zhu, Y., Liu, R., Zhang, H., Liu, S., Zhang, Z., Yu, F. H., & Gregoire, T. G. (2023). Post-flooding disturbance recovery promotes carbon capture in riparian zones. Biogeosciences, 20(7), 1357-1370.

14) Leifeld, J., & Menichetti, L. (2018). The underappreciated potential of peatlands in global climate change mitigation strategies. Nature communications, 9(1), 1071.

15) Sanders, D., & Frago, E. (2024). Ecosystem engineers shape ecological network structure and stability: A framework and literature review. Functional Ecology, 38(8), 1683-1696.

16) Rincon-Madroñero, M., Sánchez-Zapata, J. A., Barber, X., & Barbosa, J. M. (2024). Long-term vegetation responses to climate depend on the distinctive roles of rewilding and traditional grazing systems. Landscape Ecology, 39(1), 1.

17) Holdo, R. M., Holt, R. D., & Fryxell, J. M. (2009). Grazers, browsers, and fire influence the extent and spatial pattern of tree cover in the Serengeti. Ecological Applications, 19(1), 95-109.

18) Costanza, J. K., & Terando, A. J. (2019). Landscape connectivity planning for adaptation to future climate and land-use change. Current Landscape Ecology Reports, 4, 1-13.

19) Mawdsley, J. R., O’MALLEY, R. O. B. I. N., & Ojima, D. S. (2009). A review of climate-change adaptation strategies for wildlife management and biodiversity conservation. Conservation Biology, 23(5), 1080-1089.

20) Kremer, A., Ronce, O., Robledo-Arnuncio, J. J., Guillaume, F., Bohrer, G., Nathan, R., … & Schueler, S. (2012). Long-distance gene flow and adaptation of forest trees to rapid climate change. Ecology letters, 15(4), 378-392.

21) Steel, Z. L., Fogg, A. M., Burnett, R., Roberts, L. J., & Safford, H. D. (2022). When bigger isn’t better—Implications of large high-severity wildfire patches for avian diversity and community composition. Diversity and Distributions, 28(3), 439-453.

22) Skidmore, P., & Wheaton, J. (2022). Riverscapes as natural infrastructure: Meeting challenges of climate adaptation and ecosystem restoration. Anthropocene, 38, 100334.

23) Viljur, M. L., Abella, S. R., Adámek, M., Alencar, J. B. R., Barber, N. A., Beudert, B., … & Thorn, S. (2022). The effect of natural disturbances on forest biodiversity: an ecological synthesis. Biological Reviews, 97(5), 1930-1947.

24) Methorst, J., Bonn, A., Marselle, M., Böhning-Gaese, K., & Rehdanz, K. (2021). Species richness is positively related to mental health–a study for Germany. Landscape and Urban Planning, 211, 104084.

25) Massenberg, J. R., Schiller, J., & Schröter-Schlaack, C. (2023). Towards a holistic approach to rewilding in cultural landscapes. People and Nature, 5(1), 45-56.

26) Faure, E., Levrel, H., & Quétier, F. (2024). Economics of rewilding. Ambio, 53(9), 1367–1382. https://doi.org/10.1007/s13280-024-02019-2

27) Nowak-Olejnik, A., & Mocior, E. (2022). Provisioning ecosystem services of wild plants collected from seminatural habitats: A basis for sustainable livelihood and multifunctional landscape conservation. Mountain Research and Development, 42(1), R11-R19.

28) Wohl, E., Angermeier, P. L., Bledsoe, B., Kondolf, G. M., MacDonnell, L., Merritt, D. M., … & Tarboton, D. (2005). River restoration. Water Resources Research, 41(10).

29) Woodcock, B. A., Bullock, J. M., McCracken, M., Chapman, R. E., Ball, S. L., Edwards, M. E., … & Pywell, R. F. (2016). Spill-over of pest control and pollination services into arable crops. Agriculture, Ecosystems & Environment, 231, 15-23.

30) Paudel, P. K., Dhakal, S., & Sharma, S. (2024). Pathways of ecosystem-based disaster risk reduction: A global review of empirical evidence. Science of The Total Environment, 172721.

31) Basak, S. M., Hossain, M. S., Tusznio, J., & Grodzińska-Jurczak, M. (2021). Social benefits of river restoration from ecosystem services perspective: A systematic review. Environmental Science & Policy, 124, 90-100.

32) Gundersen, V., Stange, E. E., Kaltenborn, B. P., & Vistad, O. I. (2017). Public visual preferences for dead wood in natural boreal forests: The effects of added information. Landscape and Urban Planning, 158, 12-24.

33) Braczkowski, A. R., O’Bryan, C. J., Lessmann, C., Rondinini, C., Crysell, A. P., Gilbert, S., … & Biggs, D. (2023). The unequal burden of human-wildlife conflict. Communications Biology, 6(1), 182.

34) Kremen, C. (2015). Reframing the land-sparing/land-sharing debate for biodiversity conservation. Annals of the New York Academy of Sciences, 1355(1), 52-76.

35) Mishra, V., Ganguly, A. R., Nijssen, B., & Lettenmaier, D. P. (2015). Changes in observed climate extremes in global urban areas. Environmental Research Letters, 10(2), 024005.

36) Salis, M., Del Giudice, L., Jahdi, R., Alcasena-Urdiroz, F., Scarpa, C., Pellizzaro, G., … & Arca, B. (2022). Spatial patterns and intensity of land abandonment drive wildfire hazard and likelihood in Mediterranean agropastoral areas. Land, 11(11), 1942.

37) Dixon, S. J., Sear, D. A., Odoni, N. A., Sykes, T., & Lane, S. N. (2016). The effects of river restoration on catchment scale flood risk and flood hydrology. Earth Surface Processes and Landforms, 41(7), 997-1008.

38) Dale, V. H., Joyce, L. A., McNulty, S., Neilson, R. P., Ayres, M. P., Flannigan, M. D., … & Wotton, B. M. (2001). Climate change and forest disturbances: climate change can affect forests by altering the frequency, intensity, duration, and timing of fire, drought, introduced species, insect and pathogen outbreaks, hurricanes, windstorms, ice storms, or landslides. BioScience, 51(9), 723-734.

## Notes

**Competing interest statement**: The authors hereby declare that they have no conflict of interest.

### Competing Interest Statement

The authors have declared no competing interest.

## REFERENCES

Álvares, F., Blanco, J., Salvatori, V., Pimenta, V., Barroso, I., & Ribeiro, S. (2014). IBERIAN PILOT ACTION: Best practices to reduce wolf predation on free-ranging cattle in Portugal and Spain. Exploring traditional husbandry methods to reduce wolf predation on free-ranging cattle in Portugal and Spain. Final Report.

Bakker, E. S., & Svenning, J.-C. (2018). Trophic rewilding: Impact on ecosystems under global change. Philosophical Transactions of the Royal Society B: Biological Sciences, 373(1761), 20170432. 10.1098/rstb.2017.0432

Bastazini, V. A. G., Galiana, N., Hillebrand, H., Estiarte, M., Ogaya, R., Peñuelas, J., Sommer, U., & Montoya, J. M. (2021). The impact of climate warming on species diversity across scales: Lessons from experimental meta-ecosystems. Global Ecology and Biogeography, 30(7), 1545–1554. 10.1111/geb.13308

Bell, S. M., Barriocanal, C., Terrer, C., & Rosell-Melé, A. (2020). Management opportunities for soil carbon sequestration following agricultural land abandonment. Environmental Science & Policy, 108, 104–111. 10.1016/j.envsci.2020.03.018

Bennett, E. M., Cramer, W., Begossi, A., Cundill, G., Díaz, S., Egoh, B. N., Geijzendorffer, I. R., Krug, C. B., Lavorel, S., Lazos, E., Lebel, L., Martín-López, B., Meyfroidt, P., Mooney, H. A., Nel, J. L., Pascual, U., Payet, K., Harguindeguy, N. P., Peterson, G. D., … Woodward, G. (2015). Linking biodiversity, ecosystem services, and human well-being: Three challenges for designing research for sustainability. Current Opinion in Environmental Sustainability, 14, 76–85. 10.1016/j.cosust.2015.03.007

Bergin, M. D., Pedersen, R. Ø., Jensen, M., & Svenning, J.-C. (2024). Mapping rewilding potential – A systematic approach to prioritise areas for rewilding in human-dominated regions. Journal for Nature Conservation, 77, 126536. 10.1016/j.jnc.2023.126536

Blois, J. L., Zarnetske, P. L., Fitzpatrick, M. C., & Finnegan, S. (2013). Climate Change and the Past, Present, and Future of Biotic Interactions. Science, 341(6145), 499–504. 10.1126/science.1237184

Boonman, C. C. F., Huijbregts, M. A. J., Benítez-López, A., Schipper, A. M., Thuiller, W., & Santini, L. (2022). Trait-based projections of climate change effects on global biome distributions. Diversity and Distributions, 28(1), 25–37. 10.1111/ddi.13431

Bratman, G. N., Anderson, C. B., Berman, M. G., Cochran, B., de Vries, S., Flanders, J., Folke, C., Frumkin, H., Gross, J. J., Hartig, T., Kahn, P. H., Kuo, M., Lawler, J. J., Levin, P. S., Lindahl, T., Meyer-Lindenberg, A., Mitchell, R., Ouyang, Z., Roe, J., … Daily, G. C. (2019). Nature and mental health: An ecosystem service perspective. Science Advances, 5(7), eaax0903. 10.1126/sciadv.aax0903

Browning, E., Christie, M., Czajkowski, M., Chalak, A., Drummond, R., Hanley, N., Jones, K. E., Kuyer, J., & Provins, A. (2024). Valuing the economic benefits of species recovery programmes. People and Nature, 6(2), 894–905. 10.1002/pan3.10626

Burak, M. K., Ferraro, K. M., Orrick, K. D., Sommer, N. R., Ellis-Soto, D., & Schmitz, O. J. (2024). Context matters when rewilding for climate change. People and Nature, 6(2), 507–518. 10.1002/pan3.10609

Burnet, J. E., Ribeiro, D., & Liu, W. (2021). Transition and Transformation of a Rural Landscape: Abandonment and Rewilding. Sustainability, 13(9), 1–14.

Bustamante, M. M. C., Silva, J. S., Scariot, A., Sampaio, A. B., Mascia, D. L., Garcia, E., Sano, E., Fernandes, G. W., Durigan, G., Roitman, I., Figueiredo, I., Rodrigues, R. R., Pillar, V. D., de Oliveira, A. O., Malhado, A. C., Alencar, A., Vendramini, A., Padovezi, A., Carrascosa, H., … Nobre, C. (2019). Ecological restoration as a strategy for mitigating and adapting to climate change: Lessons and challenges from Brazil. Mitigation and Adaptation Strategies for Global Change, 24(7), 1249–1270. 10.1007/s11027-018-9837-5

Carroll, C., & Noss, R. F. (2021). Rewilding in the face of climate change. Conservation Biology, 35(1), 155–167. 10.1111/cobi.13531

Carver, S., Convery, I., Hawkins, S., Beyers, R., Eagle, A., Kun, Z., Van Maanen, E., Cao, Y., Fisher, M., Edwards, S. R., Nelson, C., Gann, G. D., Shurter, S., Aguilar, K., Andrade, A., Ripple, W. J., Davis, J., Sinclair, A., Bekoff, M., … Soulé, M. (2021). Guiding principles for rewilding. Conservation Biology, 35(6), 1882–1893. 10.1111/cobi.13730

Castillo, C. P., Kavalov, B., Barranco, R. R., Diogo, V., Jacobs-Crisioni, C., Silva, F. B. e, Baranzelli, C., & Lavalle, C. (2018). Territorial Fact and Trends in the EU Rural Areas within 2015-2030. JRC Research Reports, Article JRC114016. https://ideas.repec.org//p/ipt/iptwpa/jrc114016.html

Charmantier, A., McCleery, R. H., Cole, L. R., Perrins, C., Kruuk, L. E. B., & Sheldon, B. C. (2008). Adaptive Phenotypic Plasticity in Response to Climate Change in a Wild Bird Population. Science, 320(5877), 800–803. 10.1126/science.1157174

Convery, I., Carver, S., Beyers, R., Hawkins, S., Fallon, J., Derham, T., Hertel, S., Lyons, K., Locquet, A., Engel, M., Cao, Y., & Kun, Z. (2025). Editorial: Rewilding in practice. Frontiers in Conservation Science, 6. 10.3389/fcosc.2025.1561801

Cook-Patton, S. C., Drever, C. R., Griscom, B. W., Hamrick, K., Hardman, H., Kroeger, T., Pacheco, P., Raghav, S., Stevenson, M., Webb, C., Yeo, S., & Ellis, P. W. (2021). Protect, manage and then restore lands for climate mitigation. Nature Climate Change, 11(12), 1027–1034. 10.1038/s41558-021-01198-0

Cromsigt, J. P. G. M., Te Beest, M., Kerley, G. I. H., Landman, M., Le Roux, E., & Smith, F. A. (2018). Trophic rewilding as a climate change mitigation strategy? Philosophical Transactions of the Royal Society B: Biological Sciences, 373(1761), 20170440. 10.1098/rstb.2017.0440

Di Sacco, A., Hardwick, K. A., Blakesley, D., Brancalion, P. H. S., Breman, E., Cecilio Rebola, L., Chomba, S., Dixon, K., Elliott, S., Ruyonga, G., Shaw, K., Smith, P., Smith, R. J., & Antonelli, A. (2021). Ten golden rules for reforestation to optimize carbon sequestration, biodiversity recovery and livelihood benefits. Global Change Biology, 27(7), 1328–1348. 10.1111/gcb.15498

Dunn-Capper, R., Giergiczny, M., Fernández, N., Marder, F., & Pereira, H. M. (2024). Public preference for the rewilding framework: A choice experiment in the Oder Delta. People and Nature, 6(2), 610–626. 10.1002/pan3.10582

Faure, E., Levrel, H., & Quétier, F. (2024). Economics of rewilding. Ambio, 53(9), 1367–1382. 10.1007/s13280-024-02019-2

Fernández, N., Ferrier, S., Navarro, L. M., & Pereira, H. M. (2020). Essential Biodiversity Variables: Integrating In-Situ Observations and Remote Sensing Through Modeling. In J. Cavender-Bares, J. A. Gamon, & P. A. Townsend (Eds.), Remote Sensing of Plant Biodiversity (pp. 485–501). Springer International Publishing. 10.1007/978-3-030-33157-3_18

Friedman, K., Bridgewater, P., Agostini, V., Agardy, T., Arico, S., Biermann, F., Brown, K., Cresswell, I. D., Ellis, E. C., Failler, P., Kim, R. E., Pratt, C., Rice, J., Rivera, V. S., & Teneva, L. (2022). The CBD Post-2020 biodiversity framework: People’s place within the rest of nature. People and Nature, 4(6), 1475–1484. 10.1002/pan3.10403

Fuhlendorf, S. D., Engle, D. M., Kerby, J., & Hamilton, R. (2009). Pyric Herbivory: Rewilding Landscapes through the Recoupling of Fire and Grazing. Conservation Biology, 23(3), 588–598. 10.1111/j.1523-1739.2008.01139.x

García-Ruiz, J. M., Lasanta, T., Nadal-Romero, E., Lana-Renault, N., & Álvarez-Farizo, B. (2020). Rewilding and restoring cultural landscapes in Mediterranean mountains: Opportunities and challenges. Land Use Policy, 99, 104850. 10.1016/j.landusepol.2020.104850

Garrido, P., Mårell, A., Öckinger, E., Skarin, A., Jansson, A., & Thulin, C.-G. (2019). Experimental rewilding enhances grassland functional composition and pollinator habitat use. Journal of Applied Ecology, 56(4), 946–955. 10.1111/1365-2664.13338

Giergiczny, M., Czajkowski, M., Żylicz, T., & Angelstam, P. (2015). Choice experiment assessment of public preferences for forest structural attributes. Ecological Economics, 119, 8–23. 10.1016/j.ecolecon.2015.07.032

Giergiczny, M., Swenson, J. E., Zedrosser, A., & Selva, N. (2022). Large carnivores and naturalness affect forest recreational value. Scientific Reports, 12(1), 13692. 10.1038/s41598-022-17862-0

Gilbert, M., Nicolas, G., Cinardi, G., Van Boeckel, T. P., Vanwambeke, S. O., Wint, G. R. W., & Robinson, T. P. (2018). Global distribution data for cattle, buffaloes, horses, sheep, goats, pigs, chickens and ducks in 2010. Scientific Data, 5(1), 180227. 10.1038/sdata.2018.227

Griscom, B. W., Adams, J., Ellis, P. W., Houghton, R. A., Lomax, G., Miteva, D. A., Schlesinger, W. H., Shoch, D., Siikamäki, J. V., Smith, P., Woodbury, P., Zganjar, C., Blackman, A., Campari, J., Conant, R. T., Delgado, C., Elias, P., Gopalakrishna, T., Hamsik, M. R., … Fargione, J. (2017). Natural climate solutions. Proceedings of the National Academy of Sciences, 114(44), 11645–11650. 10.1073/pnas.1710465114

Hall, C. M. (2019). Tourism and rewilding: An introduction – definition, issues and review. Journal of Ecotourism, 18(4), 297–308. 10.1080/14724049.2019.1689988

Harenda, K. M., Lamentowicz, M., Samson, M., & Chojnicki, B. H. (2018). The Role of Peatlands and Their Carbon Storage Function in the Context of Climate Change. In T. Zielinski, I. Sagan, & W. Surosz (Eds.), Interdisciplinary Approaches for Sustainable Development Goals: Economic Growth, Social Inclusion and Environmental Protection (pp. 169–187). Springer International Publishing. 10.1007/978-3-319-71788-3_12

Harvey, G. L., Hartley, A. T., Henshaw, A. J., Khan, Z., Clarke, S. J., Sandom, C. J., England, J., King, S., & Venn, O. (2024). The role of rewilding in mitigating hydrological extremes: State of the evidence. WIREs Water, 11(3), e1710. 10.1002/wat2.1710

Harvey, G. L., & Henshaw, A. J. (2023). Rewilding and the water cycle. WIREs Water, 10(6), e1686. 10.1002/wat2.1686

Hering, D., Schürings, C., Wenskus, F., Blackstock, K., Borja, A., Birk, S., Bullock, C., Carvalho, L., Dagher-Kharrat, M. B., Lakner, S., Lovrić, N., McGuinness, S., Nabuurs, G.-J., Sánchez-Arcilla, A., Settele, J., & Pe’er, G. (2023). Securing success for the Nature Restoration Law. Science, 382(6676), 1248–1250. 10.1126/science.adk1658

Hermoso, V., Carvalho, S. B., Giakoumi, S., Goldsborough, D., Katsanevakis, S., Leontiou, S., Markantonatou, V., Rumes, B., Vogiatzakis, I. N., & Yates, K. L. (2022). The EU Biodiversity Strategy for 2030: Opportunities and challenges on the path towards biodiversity recovery. Environmental Science & Policy, 127, 263–271. 10.1016/j.envsci.2021.10.028

Hoffmann, A. A., & Sgrò, C. M. (2011). Climate change and evolutionary adaptation. Nature, 470(7335), 479–485. 10.1038/nature09670

Holdo, R. M., Holt, R. D., & Fryxell, J. M. (2009). Grazers, browsers, and fire influence the extent and spatial pattern of tree cover in the Serengeti. Ecological Applications, 19(1), 95–109. 10.1890/07-1954.1

IPBES. (2019). Global assessment report on biodiversity and ecosystem services of the Intergovernmental Science-Policy Platform on Biodiversity and Ecosystem Services (Version 1). Zenodo. 10.5281/ZENODO.5657041

Jones, M. W., Abatzoglou, J. T., Veraverbeke, S., Andela, N., Lasslop, G., Forkel, M., Smith, A. J. P., Burton, C., Betts, R. A., van der Werf, G. R., Sitch, S., Canadell, J. G., Santín, C., Kolden, C., Doerr, S. H., & Le Quéré, C. (2022). Global and Regional Trends and Drivers of Fire Under Climate Change. Reviews of Geophysics, 60(3), e2020RG000726. 10.1029/2020RG000726

Keenleyside, C., & Tucker, G. (2010). Farmland abandonment in the EU: An assessment of trends and prospects. London: WWF and IEEP (November.

Koninx, F. (2019). Ecotourism and rewilding: The case of Swedish Lapland. Journal of Ecotourism, 18(4), 332–347. 10.1080/14724049.2018.1538227

Lal, R. (2015). Restoring Soil Quality to Mitigate Soil Degradation. Sustainability, 7(5), Article 5. 10.3390/su7055875

Lam, D. P. M., Hinz, E., Lang, D. J., Tengö, M., Wehrden, H. V., & Martín-López, B. (2020). Indigenous and local knowledge in sustainability transformations research: A literature review. Ecology and Society, 25(1), art3. 10.5751/ES-11305-250103

Levers, C., Schneider, M., Prishchepov, A. V., Estel, S., & Kuemmerle, T. (2018). Spatial variation in determinants of agricultural land abandonment in Europe. Science of The Total Environment, 644, 95–111. 10.1016/j.scitotenv.2018.06.326

Lewis, S. L., Wheeler, C. E., Mitchard, E. T. A., & Koch, A. (2019). Restoring natural forests is the best way to remove atmospheric carbon. Nature, 568(7750), 25–28. 10.1038/d41586-019-01026-8

Malhi, Y., Doughty, C. E., Galetti, M., Smith, F. A., Svenning, J.-C., & Terborgh, J. W. (2016). Megafauna and ecosystem function from the Pleistocene to the Anthropocene. Proceedings of the National Academy of Sciences of the United States of America, 113(4), 838–846. 10.1073/pnas.1502540113

Malhi, Y., Lander, T., Le Roux, E., Stevens, N., Macias-Fauria, M., Wedding, L., Girardin, C., Kristensen, J., Sandom, C., Evans, T., Svenning, J.-C., & Canney, S. (2022). The role of large wild animals in climate change mitigation and adaptation. Current Biology, 32. 10.1016/j.cub.2022.01.041

Marjakangas, E.-L., Genes, L., Pires, M. M., Fernandez, F. A. S., de Lima, R. A. F., de Oliveira, A. A., Ovaskainen, O., Pires, A. S., Prado, P. I., & Galetti, M. (2018). Estimating interaction credit for trophic rewilding in tropical forests. Philosophical Transactions of the Royal Society B: Biological Sciences, 373(1761), 20170435. 10.1098/rstb.2017.0435

Marvin, D. C., Sleeter, B. M., Cameron, D. R., Nelson, E., & Plantinga, A. J. (2023). Natural climate solutions provide robust carbon mitigation capacity under future climate change scenarios. Scientific Reports, 13(1), 19008. 10.1038/s41598-023-43118-6

Massenberg, J. R., Schiller, J., & Schröter-Schlaack, C. (2023). Towards a holistic approach to rewilding in cultural landscapes. People and Nature, 5(1), 45–56. 10.1002/pan3.10426

McVittie, A., Cole, L., Wreford, A., Sgobbi, A., & Yordi, B. (2018). Ecosystem-based solutions for disaster risk reduction: Lessons from European applications of ecosystem-based adaptation measures. International Journal of Disaster Risk Reduction, 32, 42–54. 10.1016/j.ijdrr.2017.12.014

Merz, E., Saberski, E., Gilarranz, L. J., Isles, P. D. F., Sugihara, G., Berger, C., & Pomati, F. (2023). Disruption of ecological networks in lakes by climate change and nutrient fluctuations. Nature Climate Change, 13(4), 389–396. 10.1038/s41558-023-01615-6

Mittelman, P., Landim, A. R., Genes, L., Assis, A. P. A., Starling-Manne, C., Leonardo, P. V., Fernandez, F. A. S., Guimarães Jr, P. R., & Pires, A. S. (2022). Trophic rewilding benefits a tropical community through direct and indirect network effects. Ecography, 2022(4). 10.1111/ecog.05838

Mokany, K., Thomson, J. J., Lynch, A. J. J., Jordan, G. J., & Ferrier, S. (2015). Linking changes in community composition and function under climate change. Ecological Applications, 25(8), 2132–2141. 10.1890/14-2384.1

Morelli, T. L., Daly, C., Dobrowski, S. Z., Dulen, D. M., Ebersole, J. L., Jackson, S. T., Lundquist, J. D., Millar, C. I., Maher, S. P., Monahan, W. B., Nydick, K. R., Redmond, K. T., Sawyer, S. C., Stock, S., & Beissinger, S. R. (2016). Managing Climate Change Refugia for Climate Adaptation. PLOS ONE, 11(8), e0159909. 10.1371/journal.pone.0159909

Munang, R., Thiaw, I., Alverson, K., Liu, J., & Han, Z. (2013). The role of ecosystem services in climate change adaptation and disaster risk reduction. Current Opinion in Environmental Sustainability, 5(1), 47–52. 10.1016/j.cosust.2013.02.002

Nayak, N., Mehrotra, R., & Mehrotra, S. (2022). Carbon biosequestration strategies: A review. Carbon Capture Science & Technology, 4, 100065. 10.1016/j.ccst.2022.100065

Palmero-Iniesta, M., Espelta, J. M., Padial-Iglesias, M., Gonzàlez-Guerrero, Ò., Pesquer, L., Domingo-Marimon, C., Ninyerola, M., Pons, X., & Pino, J. (2021). The Role of Recent (1985– 2014) Patterns of Land Abandonment and Environmental Factors in the Establishment and Growth of Secondary Forests in the Iberian Peninsula. Land, 10(8), Article 8. 10.3390/land10080817

Paudel, P. K., Dhakal, S., & Sharma, S. (2024). Pathways of ecosystem-based disaster risk reduction: A global review of empirical evidence. Science of The Total Environment, 929, 172721. 10.1016/j.scitotenv.2024.172721

Pearce, E. A., Mazier, F., Davison, C. W., Baines, O., Czyżewski, S., Fyfe, R., Bińka, K., Boreham, S., de Beaulieu, J.-L., Gao, C., Granoszewski, W., Hrynowiecka, A., Malkiewicz, M., Mighall, T., Noryśkiewicz, B., Pidek, I. A., Strahl, J., Winter, H., & Svenning, J.-C. (2025). Beyond the closed-forest paradigm: Cross-scale vegetation structure in temperate Europe before the late-Quaternary megafauna extinctions. Earth History and Biodiversity, 3, 100022. 10.1016/j.hisbio.2025.100022

Pecl, G. T., Araújo, M. B., Bell, J. D., Blanchard, J., Bonebrake, T. C., Chen, I.-C., Clark, T. D., Colwell, R. K., Danielsen, F., Evengård, B., Falconi, L., Ferrier, S., Frusher, S., Garcia, R. A., Griffis, R. B., Hobday, A. J., Janion-Scheepers, C., Jarzyna, M. A., Jennings, S., … Williams, S. E. (2017). Biodiversity redistribution under climate change: Impacts on ecosystems and human well-being. Science, 355(6332), eaai9214. 10.1126/science.aai9214

Pellis, A. (2019). Reality effects of conflict avoidance in rewilding and ecotourism practices – the case of Western Iberia. Journal of Ecotourism, 18(4), 316–331. 10.1080/14724049.2019.1579824

Pereira, H. M., Ferrier, S., Walters, M., Geller, G. N., Jongman, R. H. G., Scholes, R. J., Bruford, M. W., Brummitt, N., Butchart, S. H. M., Cardoso, A. C., Coops, N. C., Dulloo, E., Faith, D. P., Freyhof, J., Gregory, R. D., Heip, C., Höft, R., Hurtt, G., Jetz, W., … Wegmann, M. (2013). Essential Biodiversity Variables. Science, 339(6117), 277–278. 10.1126/science.1229931

Pereira, H. M., & Navarro, L. M. (Eds.). (2015). Rewilding European Landscapes. Springer International Publishing. 10.1007/978-3-319-12039-3

Perino, A., Pereira, H. M., Navarro, L. M., Fernández, N., Bullock, J. M., Ceaușu, S., Cortés-Avizanda, A., Van Klink, R., Kuemmerle, T., Lomba, A., Pe’er, G., Plieninger, T., Rey Benayas, J. M., Sandom, C. J., Svenning, J.-C., & Wheeler, H. C. (2019). Rewilding complex ecosystems. Science, 364(6438), eaav5570. 10.1126/science.aav5570

Pettorelli, N., Durant, S. M., & Du Toit, J. T. (Eds.). (2019). Rewilding (1st ed.). Cambridge University Press. 10.1017/9781108560962

Pimenta, V., Barroso, I., Boitani, L., & Beja, P. (2017). Wolf predation on cattle in Portugal: Assessing the effects of husbandry systems. Biological Conservation, 207, 17–26. 10.1016/j.biocon.2017.01.008

Pimenta, V., Barroso, I., Boitani, L., & Beja, P. (2018). Risks *a la carte*: Modelling the occurrence and intensity of wolf predation on multiple livestock species. Biological Conservation, 228, 331–342. 10.1016/j.biocon.2018.11.008

Plutzar, C., Kroisleitner, C., Haberl, H., Fetzel, T., Bulgheroni, C., Beringer, T., Hostert, P., Kastner, T., Kuemmerle, T., Lauk, C., Levers, C., Lindner, M., Moser, D., Müller, D., Niedertscheider, M., Paracchini, M. L., Schaphoff, S., Verburg, P. H., Verkerk, P. J., & Erb, K.-H. (2016). Changes in the spatial patterns of human appropriation of net primary production (HANPP) in Europe 1990–2006. Regional Environmental Change, 16(5), 1225–1238. 10.1007/s10113-015-0820-3

Pörtner, H.-O., Scholes, R. J., Arneth, A., Barnes, D. K. A., Burrows, M. T., Diamond, S. E., Duarte, C. M., Kiessling, W., Leadley, P., Managi, S., McElwee, P., Midgley, G., Ngo, H. T., Obura, D., Pascual, U., Sankaran, M., Shin, Y. J., & Val, A. L. (2023). Overcoming the coupled climate and biodiversity crises and their societal impacts. Science, 380(6642), eabl4881. 10.1126/science.abl4881

Rabin, S. S., Gérard, F. N., & Arneth, A. (2022). The influence of thinning and prescribed burning on future forest fires in fire-prone regions of Europe. Environmental Research Letters, 17(5), 055010. 10.1088/1748-9326/ac6312

Radchuk, V., Reed, T., Teplitsky, C., van de Pol, M., Charmantier, A., Hassall, C., Adamík, P., Adriaensen, F., Ahola, M. P., Arcese, P., Miguel Avilés, J., Balbontin, J., Berg, K. S., Borras, A., Burthe, S., Clobert, J., Dehnhard, N., de Lope, F., Dhondt, A. A., … Kramer-Schadt, S. (2019). Adaptive responses of animals to climate change are most likely insufficient. Nature Communications, 10(1), 3109. 10.1038/s41467-019-10924-4

Rey Benayas, J. M., Martins, A., Nicolau, J. M., & Schulz, J. J. (2007). Abandonment of agricultural land: An overview of drivers and consequences. CABI Reviews, 2007, 14 pp. 10.1079/PAVSNNR20072057

Root-Bernstein, M., Gooden, J., & Boyes, A. (2018). Rewilding in practice: Projects and policy. Geoforum, 97, 292–304. 10.1016/j.geoforum.2018.09.017

Sandom, C. J., Middleton, O., Lundgren, E., Rowan, J., Schowanek, S. D., Svenning, J.-C., & Faurby, S. (2020). Trophic rewilding presents regionally specific opportunities for mitigating climate change. Philosophical Transactions of the Royal Society B: Biological Sciences, 375(1794), 20190125. 10.1098/rstb.2019.0125

Scarano, F. R. (2017). Ecosystem-based adaptation to climate change: Concept, scalability and a role for conservation science. Perspectives in Ecology and Conservation, 15(2), 65–73. 10.1016/j.pecon.2017.05.003

Schmitz, O. J., Sylvén, M., Atwood, T. B., Bakker, E. S., Berzaghi, F., Brodie, J. F., Cromsigt, J. P. G. M., Davies, A. B., Leroux, S. J., Schepers, F. J., Smith, F. A., Stark, S., Svenning, J.-C., Tilker, A., & Ylänne, H. (2023). Trophic rewilding can expand natural climate solutions. Nature Climate Change, 13(4), 324–333. 10.1038/s41558-023-01631-6

Schou, J. S., Bladt, J., Ejrnæs, R., Thomsen, M. N., Vedel, S. E., & Fløjgaard, C. (2021). Economic assessment of rewilding versus agri-environmental nature management. Ambio, 50(5), 1047–1057. 10.1007/s13280-020-01423-8

Selwyn, M., Lázaro-González, A., Lloret, F., Rey Benayas, J. M., Hampe, A., Brotons, L., Pino, J., & Espelta, J. M. (2025). Quantifying the impacts of rewilding on ecosystem resilience to disturbances: A global meta-analysis. Journal of Environmental Management, 375, 124360. 10.1016/j.jenvman.2025.124360

Seneviratne, S. I., Zhang, X., Adnan, M., Badi, W., Dereczynski, C., Luca, A. D., Ghosh, S., Iskandar, I., Kossin, J., Lewis, S., Otto, F., Pinto, I., Satoh, M., Vicente-Serrano, S. M., Wehner, M., Zhou, B., & Allan, R. (2021). Weather and climate extreme events in a changing climate (V. P. Masson-Delmotte, A. Zhai, S. L. Pirani, & C. Connors, Eds.; pp. 1513–1766). Cambridge University Press. https://centaur.reading.ac.uk/101846/

Siikamäki, J., Sanchirico, J. N., Jardine, S., McLaughlin, D., & Morris, D. (2013). Blue Carbon: Coastal Ecosystems, Their Carbon Storage, and Potential for Reducing Emissions. Environment: Science and Policy for Sustainable Development, 55(6), 14–29. 10.1080/00139157.2013.843981

Sonntag, S., & Fourcade, Y. (2022). Where will species on the move go? Insights from climate connectivity modelling across European terrestrial habitats. Journal for Nature Conservation, 66, 126139. 10.1016/j.jnc.2022.126139

Sowińska-Świerkosz, B., & García, J. (2022). What are Nature-based solutions (NBS)? Setting core ideas for concept clarification. Nature-Based Solutions, 2, 100009. 10.1016/j.nbsj.2022.100009

Stark, G., & Galetti, M. (2024). Rewilding in cold blood: Restoring functionality in degraded ecosystems using herbivorous reptiles. Global Ecology and Conservation, 50, e02834. 10.1016/j.gecco.2024.e02834

Stark, G., Ma, L., Zeng, Z.-G., Du, W.-G., & Levy, O. (2023). Cool shade and not-so-cool shade: How habitat loss may accelerate thermal stress under current and future climate. Global Change Biology, 29(22), 6201–6216. 10.1111/gcb.16802

Stark, G., & Schwarz, R. (2024). Rewilding a vanishing taxon – Restoring aquatic ecosystems using amphibians. Biological Conservation, 292, 110559. 10.1016/j.biocon.2024.110559

Stephens, T. (2023). The Kunming–Montreal Global Biodiversity Framework. International Legal Materials, 62(5), 868–887. 10.1017/ilm.2023.16

Suárez-Ronay, S. H., Medina-Villar, S., & Corona, M. E. P. (2024). The role of fire in the germination of invasive plants in Mediterranean environments: A meta-analysis. Forest Ecology and Management, 569, 122168. 10.1016/j.foreco.2024.122168

Summers, J. K., Smith, L. M., Case, J. L., & Linthurst, R. A. (2012). A Review of the Elements of Human Well-Being with an Emphasis on the Contribution of Ecosystem Services. AMBIO, 41(4), 327–340. 10.1007/s13280-012-0256-7

Suziedelyte Visockiene, J., Tumeliene, E., & Maliene, V. (2019). Analysis and identification of abandoned agricultural land using remote sensing methodology. Land Use Policy, 82, 709–715. 10.1016/j.landusepol.2019.01.013

Svenning, J.-C. (2020). Rewilding should be central to global restoration efforts. One Earth, 3(6), 657–660. 10.1016/j.oneear.2020.11.014

Svenning, J.-C., Buitenwerf, R., & Roux, E. L. (2024). Trophic rewilding as a restoration approach under emerging novel biosphere conditions. Current Biology, 34(9), R435–R451. 10.1016/j.cub.2024.02.044

Svenning, J.-C., Pedersen, P. B. M., Donlan, C. J., Ejrnæs, R., Faurby, S., Galetti, M., Hansen, D. M., Sandel, B., Sandom, C. J., Terborgh, J. W., & Vera, F. W. M. (2016). Science for a wilder Anthropocene: Synthesis and future directions for trophic rewilding research. Proceedings of the National Academy of Sciences, 113(4), 898–906. 10.1073/pnas.1502556112

Thakur, M. P., Bakker, E. S., Veen, G. F. (Ciska), & Harvey, J. A. (2020). Climate Extremes, Rewilding, and the Role of Microhabitats. One Earth, 2(6), 506–509. 10.1016/j.oneear.2020.05.010

Thulin, C.-G., & Röcklinsberg, H. (2020). Ethical Considerations for Wildlife Reintroductions and Rewilding. Frontiers in Veterinary Science, 7, 163. 10.3389/fvets.2020.00163

Torres, A., Fernández, N., Zu Ermgassen, S., Helmer, W., Revilla, E., Saavedra, D., Perino, A., Mimet, A., Rey-Benayas, J. M., Selva, N., Schepers, F., Svenning, J.-C., & Pereira, H. M. (2018). Measuring rewilding progress. Philosophical Transactions of the Royal Society B: Biological Sciences, 373(1761), 20170433. 10.1098/rstb.2017.0433

Vieira, D. C. S., Borrelli, P., Jahanianfard, D., Benali, A., Scarpa, S., & Panagos, P. (2023). Wildfires in Europe: Burned soils require attention. Environmental Research, 217, 114936. 10.1016/j.envres.2022.114936

Villar, N., & Medici, E. P. (2021). Large wild herbivores slow down the rapid decline of plant diversity in a tropical forest biodiversity hotspot. Journal of Applied Ecology, 58(11), 2361–2370. 10.1111/1365-2664.14054

von Holle, B., Yelenik, S., & Gornish, E. S. (2020). Restoration at the landscape scale as a means of mitigation and adaptation to climate change. Current Landscape Ecology Reports, 5(3), 85–97. 10.1007/s40823-020-00056-7

Waylen, K. A., Wilkinson, M. E., Blackstock, K. L., & Bourke, M. (2024). Nature-based solutions and restoration are intertwined but not identical: Highlighting implications for societies and ecosystems. Nature-Based Solutions, 5, 100116. 10.1016/j.nbsj.2024.100116

Williams, J., Sandom, C. J., & Pettorelli, N. (2024). Active management is required to regenerate the Caledonian forest: Alladale as a case study. Ecological Solutions and Evidence, 5(1), e12315. 10.1002/2688-8319.12315

Wilson, G. R., & Edwards, M. J. (2008). Native wildlife on rangelands to minimize methane and produce lower-emission meat: Kangaroos versus livestock. Conservation Letters, 1(3), 119–128. 10.1111/j.1755-263X.2008.00023.x

Winder, M., & Schindler, D. E. (2004). Climatic effects on the phenology of lake processes. Global Change Biology, 10(11), 1844–1856. 10.1111/j.1365-2486.2004.00849.x

zu Ermgassen, S. O. S. E., McKenna, T., Gordon, J., & Willcock, S. (2018). Ecosystem service responses to rewilding: First-order estimates from 27 years of rewilding in the Scottish Highlands. International Journal of Biodiversity Science, Ecosystem Services & Management, 14(1), 165–178. 10.1080/21513732.2018.1502209

