## Supplementary materials for "TOWARDS CLIMATE-SMART REWILDING: AN INTEGRATED FRAMEWORK FOR BIODIVERSITY, CLIMATE CHANGE, AND SOCIETY"

A. Data sources for the spatial assessment for potential rewilding areas across Europe – Climate mitigation (Land abandonment & Potential Net Primary Productivity) and adaptation (Land abandonment & Climate velocity).

Spatial data for both maps was meticulously gathered from various scientific literature sources and global database products accessible via online platforms. Through the process of overlapping the maps, we were able to illustrate a workflow for delineating potential areas suitable for implementing climate-smart rewilding initiatives.

**Potential Net Primary Productivity**

The data utilized for our study on potential NPP was sourced from Plutzar et al. (2016). This dataset offers a comprehensive assessment of potential NPP for Europe across a 1-kilometer grid for the years 1990, 2000, and 2006. We used for our example the Potential NPP map calculated for 2006. Our research specifically focused on comparing the distribution of land abandonment with NPP areas. This allowed us to examine the potential impact of conserving areas with high primary productivity (e.g., expanding forests on abandoned land) to increase potential above-ground carbon sequestration. Through this analysis, we aimed to gain insights into the relationship between farmland abandonment and landscape productivity.

**Climate velocity**

To construct a climate velocity map, we calculated the rate and direction of climate change within a designated region, employing the methodology established by Loarie et al., (2009). Our analysis utilized climate data from the WorldClim database version 2.1 (Fick & Hijmans, 2017), which was released in January 2020. We incorporated two distinct datasets: one pertaining to historical climate data and the other to future climate projections. The historical climate data focused solely on the BIO1 bioclimatic variable, which represents the Annual Mean Temperature. This dataset was obtained at a spatial resolution of 30 seconds (approximately 1 km²) and covers the period from 1970 to 2000. Conversely, the future climate data also concentrated exclusively on the BIO1 variable, based on the CMIP6 downscaled future climate projections. This dataset, maintaining the same spatial resolution of 30 seconds, provides average temperature data for the interval between 2061 and 2080. The predictive model employed for these future projections was the CMCC-EMS2, utilizing the SSP245 scenario. To ascertain the temporal gradient, we first determined the total temperature difference between the historical and future periods and subsequently divided this difference by the number of years encompassed.

$$Temporal\ Gradient = \frac{t_1 - t_2}{N.of\ Years} = \frac{\Delta Temp.}{\Delta Time}$$

When applied:

$$Temporal\ Gradient = \frac{Future\ Climate - Historical\ Climate}{80}$$

The spatial gradient is based on Horn’s Methods for the slope (Horn, 1981):

| | | |
|---|---|---|
| A | B | C |
| D | E | F |
| G | H | I |

$$\frac{dz}{dx} = \frac{(c + 2f + i) - (a + 2d + g)}{8}$$

$$\frac{dz}{dy} = \frac{(g + 2h + i) - (a + 2b + c)}{8}$$

$$Slope\ (spatial\ gradient) = \sqrt{\frac{dz}{dx}^2 + \frac{dz}{dy}^2}$$

To get the climate change velocity, the temporal gradient is divided by the spatial gradient:

$$Ve = \frac{Temporal\ Gradient}{Spatial\ Gradient}$$

This computation generates the rate of temperature change across a gradient (°C km$^{-1}$) in an average year (km yr$^{-1}$).

**Land abandonment**

We used spatial data on the projections of land abandonment covering the years 2000 to 2040, based on various scenarios outlined by Ceaușu et al. (2015). In their study, they employed the Dyna-CLUE model's land-use change projections at a 1 km² resolution (Verburg & Overmars, 2009) to pinpoint areas in Europe experiencing farmland abandonment, alongside four socio-economic VOLANTE scenarios that represent different policy and management choices in the region (Ceaușu et al., 2015). Abandonment was only considered if it was predicted in at least three of these scenarios. This assessment is crucial for understanding potential environmental and socio-economic changes within the specified timeframe. By examining diverse scenarios, the goal was to capture a thorough understanding of how land abandonment might progress under varying conditions, including shifts in agricultural practices, urban development, and policy adjustments. The insights gained from these projections shed light on the future land-use change and its implications for land use and management strategies.

B. Spatial Assessment for potential rewilding areas across Europe – Large mammal expansion and human-wildlife co-existence challenges.

The social benefits and conflicts arising from our mapping are contingent upon the expanding trophic complexity that results from rewilding initiatives. Our focus is on the resurgence of large mammal species, such as wolves (*Canis lupus lupus*) and elks (*Cervus canadensis*). Their expansion into new territories could offer enhanced opportunities for wildlife observation, enjoyment, and interaction with nature for people. However, this also poses a threat to livestock (in the case of large predators), leading to potential conflicts with humans.

**Wildlife comeback**

We assessed wildlife recolonization potential using a 2018 Europe-wide large mammal distribution dataset (*Boosting Ecological Restoration for a Wilder Europe: Making the Green Deal Work for Nature*, 2020) comprising presence/absence data for 14 species. For each grid cell lacking a species' presence, we calculated the minimum distance to the nearest occupied cell (D, in km). Colonization probability was then estimated using the decay function:

$$1 \,/\, exp\,[(log\,(2)/60)\; *\; D]$$

Cells with existing species or at great distances from occupied cells had a colonization probability of 0.

Figure 3 categorizes areas based on four wildlife statuses: 1) at least one species present, no colonization; 2) at least one species present, ≥0.8 colonization probability; 3) no species present, ≥0.8 colonization probability; 4) no species present, zero colonization probability.

**Socio-economic benefits**

One example of estimating social benefits from rewilding involves identifying the areas where a specific, often charismatic or flagship species, might return and where people would likely travel to observe it. This latter element is measured through willingness-to-travel (WTT), which represents the distance people are willing to travel to see a certain species. Using survey data on the recreational value of two large carnivore species: the Brown bear (*Ursus arctos*), and the Eurasian wolf (*Canis lupus lupus*; Giergiczny et al., 2022), we established the correlation between the distance traveled to a specific location containing a large carnivore species and the percentage of individuals willing to travel that particular distance (represented by 1-CDF of the lognormal distribution with parameters $\mu$ = -250 and $\sigma$ = 100). Schematically a high percentage of people (e.g. > 90%) are willing to travel a short distance (e.g. 1km) and a small percentage of people (e.g. < 10%) are willing to travel a long distance (e.g. 100km). The JRC-GEOSTAT Population Grid 2018 was utilized to allocate the total human population that would potentially travel to each location, derived from a kernel convolution based on this correlation. In this analysis eight species were included: Brown bear (*Ursus arctos*), Eurasian wolf (*Canis lupus lupus*), Elk (*Cervus canadensis*), European bison (*Bison bonasus*), Eurasian lynx (*Lynx lynx*), Iberian lynx (*Lynx pardinus*), Alpine ibex (*Capra ibex*), and Iberian ibex (*Capra pyrenaica*). We summed up the probability of colonization of those eight species.

**Socio-economic risks**

One example of estimating the risks for society associated with rewilding involves identifying regions where species that threaten livestock, whether through predation or competition, overlap with areas where domesticated animals are present (i.e., the number of livestock in a specific grid cell). This assessment involves examining the distribution of cattle, sheep, and goats across Europe using data from the Gridded Livestock of the World (GLW) database (Gilbert et al., 2018). In this analysis, the Brown bear and European wolf are considered potential threats. We summed the probability of colonization of those two species.

**REFERENCES**


*Boosting Ecological Restoration for a Wilder Europe: Making the Green Deal Work for Nature* (with Fernández, N., Torres, A., Wolf, F., Quintero, L., & Pereira, H. M.). (2020). Martin-Luther-Universität.

Ceaușu, S., Hofmann, M., Navarro, L. M., Carver, S., Verburg, P. H., & Pereira, H. M. (2015). Mapping opportunities and challenges for rewilding in Europe. *Conservation Biology*, *29*(4), 1017–1027. https://doi.org/10.1111/cobi.12533

Fick, S. E., & Hijmans, R. J. (2017). WorldClim 2: New 1-km spatial resolution climate surfaces for global land areas. *International Journal of Climatology*, *37*(12), 4302–4315. https://doi.org/10.1002/joc.5086

Giergiczny, M., Swenson, J. E., Zedrosser, A., & Selva, N. (2022). Large carnivores and naturalness affect forest recreational value. *Scientific Reports*, *12*(1), 13692. https://doi.org/10.1038/s41598-022-17862-0

Gilbert, M., Nicolas, G., Cinardi, G., Van Boeckel, T. P., Vanwambeke, S. O., Wint, G. R. W., & Robinson, T. P. (2018). Global distribution data for cattle, buffaloes, horses, sheep, goats, pigs, chickens and ducks in 2010. *Scientific Data*, *5*(1), 180227. https://doi.org/10.1038/sdata.2018.227

Horn, B. K. P. (1981). Hill shading and the reflectance map. *Proceedings of the IEEE*, *69*(1), 14–47. Proceedings of the IEEE. https://doi.org/10.1109/PROC.1981.11918

Loarie, S. R., Duffy, P. B., Hamilton, H., Asner, G. P., Field, C. B., & Ackerly, D. D. (2009). The velocity of climate change. *Nature*, *462*(7276), 1052–1055. https://doi.org/10.1038/nature08649

Plutzar, C., Kroisleitner, C., Haberl, H., Fetzel, T., Bulgheroni, C., Beringer, T., Hostert, P., Kastner, T., Kuemmerle, T., Lauk, C., Levers, C., Lindner, M., Moser, D., Müller, D., Niedertscheider, M., Paracchini, M. L., Schaphoff, S., Verburg, P. H., Verkerk, P. J., & Erb, K.-H. (2016). Changes in the spatial patterns of human appropriation of net primary production (HANPP) in Europe 1990–2006. *Regional Environmental Change*, *16*(5), 1225–1238. https://doi.org/10.1007/s10113-015-0820-3

Verburg, P. H., & Overmars, K. P. (2009). Combining top-down and bottom-up dynamics in land use modeling: Exploring the future of abandoned farmlands in Europe with the Dyna-CLUE model. *Landscape Ecology*, *24*(9), 1167–1181. https://doi.org/10.1007/s10980-009-9355-7
